# The Evolution of Cooperation by the Hankshaw Effect

**DOI:** 10.1101/016667

**Authors:** Sarah Hammarlund, Brian D. Connelly, Katherine J. Dickinson, Benjamin Kerr

## Abstract

The evolution of cooperation—costly behavior that benefits others—faces one clear obstacle. Namely, cooperators are always at a competitive disadvantage relative to defectors, individuals that reap the same social benefits, but evade the personal cost. One solution to this problem involves genetic hitchhiking, where the allele encoding cooperative behavior becomes linked to a beneficial mutation. While traditionally seen as a passive process driven purely by chance, here we explore a more active form of hitchhiking. Specifically, we model hitchhiking in the context of adaptation to a stressful environment by cooperators and defectors with spatially limited dispersal. Under such conditions, clustered cooperators reach higher local densities, thereby experiencing more opportunities for mutations than defectors. Thus, the allele encoding cooperation has a greater probability of hitchhiking with alleles conferring stress adaptation. We label this probabilistic enhancement the “Hankshaw effect” after the character Sissy Hankshaw, whose anomalously large thumbs made her a singularly effective hitchhiker. Using an agent-based model, we demonstrate that there exists a broad set of conditions allowing the evolution of cooperation through the Hankshaw effect. We discuss the feasibility of our theoretical assumptions for natural systems, not only for the case of cooperation, but also for other costly social behaviors such as spite. The primary elements of our model, including genetic hitchhiking and population structure, have been discussed separately in previous models exploring the evolution of cooperation. However, the combination of these elements has not been appreciated as a solution to the problem of cooperation.

## Introduction

A deleterious allele can increase in frequency if it is physically linked to a beneficial allele (Maynard Smith and Haigh 1974). Such genetic hitchhiking is often viewed as a passive process—the deleterious allele becomes associated with a positively selected allele purely by chance. However, in some cases the hitchhiking allele can play an active role by increasing its probability of catching a ride. For instance, an allele that increases the genomic mutation rate may lift its chances of hitchhiking by raising the incidence of beneficial mutations, despite the deleterious mutations it also generates (de Visser 2002). If a property of an allele increases its likelihood of hitchhiking, we term this the “Hankshaw effect,” after the character Sissy Hankshaw from Tom Robbins’ novel *Even Cowgirls Get the Blues*. Hankshaw was born with oversized thumbs and uses this attribute to become a prolific hitchhiker. For Hankshaw, a trait that was initially an impairment becomes her salvation on the open road. In the same way, the cost of a deleterious allele can be offset if the allele improves its own chances of hitchhiking. Here, we explore how the Hankshaw effect can promote the evolution of one costly trait that has received a great deal of attention: cooperation.

We define cooperation as costly behavior that improves the fitness of others. For instance, the production of costly secreted enzymes by microbes can liberate critical resources or detoxify harmful substances present in the environment; and thus these exoenzymes constitute public goods (Greig and Travisano 2004, West et al. 2007a, Sandoz et al. 2007, Dugatkin et al. 2005). It is the cost of cooperative behavior that makes its evolution so problematic. Specifically, a population of cooperators is susceptible to invasion by defectors—individuals that forego the costs of cooperation but still reap its benefits (Hardin 1968, Velicer et al. 2000, Strassmann et al. 2000, Rainey and Rainey 2003, Travisano and Velicer 2004). One recently proposed solution to this subversion problem involves genetic hitchhiking (Waite and Shou 2012, Morgan et al., 2012, Asfahl et al. 2015). In these studies, bacteria or yeast that produce public goods (cooperators) compete against non-producers (defectors) in a novel environment. If a beneficial mutation happens to arise first in the cooperative strain, and if the selective advantage of this mutation outweighs the cost of cooperation, this adapted cooperator can displace defectors through genetic hitchhiking. These studies have focused on this hitchhiking process in well-mixed populations of cooperators and defectors. Under such conditions, cooperators do not have a greater chance of acquiring a beneficial mutation.

However, there are circumstances where cooperators can increase their chances of adaptation. Specifically, if the population is spatially structured (in which limited dispersal leads to clustering of like-types) the benefits of cooperation will be disproportionately experienced by cooperators, and cooperator-rich regions will reach higher densities. Consequently, cooperative lineages expanding within a structured population will experience more reproduction than defector lineages and therefore more opportunities for a beneficial mutation. Because cooperators adapt more rapidly, they are able to competitively displace defectors when the two types meet. That is, cooperator alleles hitchhike with the beneficial mutations that are more likely to occur in their presence. In structured populations, such evolution of cooperation by the Hankshaw effect can occur as long as there is room for evolutionary improvement.

There is ample opportunity for adaptation under stressful environmental conditions, as organisms experiencing such conditions are, by definition, maladapted to their environment. Evolution under harsh conditions can involve selection for mutations that allow organisms to better tolerate the stress. Stressful conditions can also thin and fragment a population. For instance, immediately after application of an antibiotic, a bacterial population can be dramatically reduced in density as only resistant types can grow. If there is considerable reduction in population size, and dispersal and migration are spatially restricted, then like-types will end up clustered together. Under such positive assortment, the benefits of cooperation are disproportionately experienced by cooperators. This clustering immediately gives cooperators a numerical advantage (Nowak 2006, Brockhurst 2007), and thereby accelerates the rate of adaptation. For example, following antibiotic exposure, bacterial survivors that cooperate (e.g., produce public goods) may have greater opportunities to compensate for the costs of resistance. Given sufficient stress adaptation, cooperators overcome the cost of cooperation and exceed the fitness of defectors, thus giving cooperators a competitive advantage. Here, we build a simulation model to explore such evolution of cooperation by the Hankshaw effect.

## Methods

In our agent-based simulation, evolution occurs in a metapopulation consisting of populations connected by limited migration. There are two types of individuals within populations: cooperators and defectors. Cooperation is costly, but increases the productivity of the population. A simulated stress thins the metapopulation at the beginning of the simulation, and then surviving lineages can acquire fitness-enhancing mutations to adapt to the stress.

### Individual genotype and fitness

The genotype of each individual is a binary string of length *L* + 1. Values (alleles) in the first *L* positions (loci) determine the individual’s level of adaptation to the stressful environment. We refer to these loci as “stress loci.” A mutation from 0 to 1 at stress locus *i* will improve individual fitness by *w*_*i*_ regardless of the allelic states of other loci (i.e., there is no epistasis). We assume that are {*w*_1_, *w*_2_, *w*_3_,…,*w*_*L*_} independent and identically distributed random variables with *w*_*i*_ ∼ unif(*w*_min_, *w*_max_). The allele at locus *L* + 1 determines whether the individual is a cooperator (allele 1) or a defector (allele 0). We refer to this last locus as the “cooperation locus.” Cooperation is costly, reducing individual fitness by *c*. Thus, if the allelic state of locus *i* is denoted *a*_*i*_ (with *a*_*i*_ ∈ {0,1}), then the fitness of an individual is:

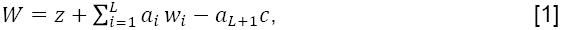

where *z* is a baseline fitness (the fitness of an individual with zeros at every locus). If there are no stress loci (*L* = 0), then the fitnesses of a cooperator and a defector are *z* - *c* and *z*, respectively.

### Overview of the metapopulation and basic simulation cycle

Simulations using this model track a single metapopulation with *N*^2^ sites arranged as an *N*×*N* bounded lattice. Each site can potentially hold a population. Simulations are run for *T* cycles and all populations cycle synchronously. Each cycle consists of population growth, mutation, migration, and dilution.

### Population growth

If *p* is the proportion of cooperators in a population at the beginning of a growth cycle, then that population grows to size *S*(*p*) over the growth cycle, where

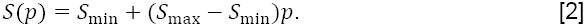

Therefore, a population consisting entirely of defectors (*p* = 0) reaches size *S*_min_, while a population of cooperators (*p* = 1) reaches a size of *S*_max_ (with *S*_max_≥*S*_min_). The function *S*(*p*) gauges the benefit of cooperation, as population size increases linearly with the proportion of cooperators. During population growth, competition among genotypes occurs. There are 2^*L*+1^ possible genotypes. Consider an arbitrary genotype *g* (with *g* ∈ {1,2,3,…,2^*L*+1^}). Let *n*_*g*_ be the number of individuals with genotype *g*, and let *w*_*g*_ be the fitness of genotype *g* (see Equation [1]). The composition of genotypes after population growth is multinomial with parameters *S*(*p*) and {*π*_1_, *π*_2_,…, π_2^*L*+1^_}, where

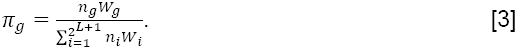

Thus, *π*_*g*_ is the probability that an individual in the population is genotype *g* after growth (such that ∑*π*_*g*_ = 1). We are therefore modeling a form of fecundity selection in an asexually reproducing population. Such selection occurs at every occupied site in the metapopulation.

### Mutation

For simplicity, mutation occurs after population growth. For each individual, every locus mutates independently. Each stress locus changes allelic state with probability *μ*_*s*_, while the cooperation locus changes allelic state with probability *μ*_*c*_. Thus, the probability that genotype *g* mutates into genotype *g*′ is given by

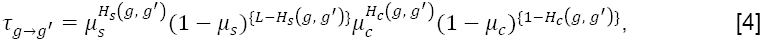

where *H*_*s*_(*g*, *g*′) and *H*_*c*_(*g*, *g*′) are the Hamming distances between genotypes *g* and *g*′ at the stress loci and cooperation locus, respectively. The Hamming distance between two genotypes is the number of loci at which those genotypes differ.

### Migration

Following mutation, individuals can migrate to new populations. For each populated site, a neighboring destination site is chosen. For a focal site in the interior of the lattice, its destination site is in its Moore neighborhood, consisting of the 8 nearest sites. The metapopulation lattice has boundaries; therefore, the destination site for a focal site on the edge or corner of the lattice is one of the nearest 5 or 3 neighboring sites, respectively. Each individual in the focal site moves to the destination site with probability *m*.

### Dilution

After migration, populations are thinned to allow for growth in the next cycle. Each individual, despite its genotype, survives dilution with probability *d*. Specifically, 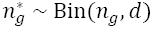, where 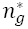 is the number of survivors for each genotype *g*.

### Stress survival and adaptation

Environmental stress has multiple effects. First, the populations undergo a bottleneck. For most runs, this thinning occurs only at the beginning of the simulation (but see the next section). This dramatic bottleneck is distinct from the mild dilution that occurs every cycle. Individuals survive the onset of the stress with probability *μ*_*t*_, which represents the likelihood of acquiring a mutation conferring stress tolerance (where *μ*_*t*_ ≪ *d*). Second, the allelic state *a*_*i*_ is set to 0 at each stress locus, as individuals are not adapted to the new stressful conditions. Third, the fitness increments tied to adaptations at each locus (*w*_*i*_) are determined as described above. All simulations begin by applying these effects to full populations initiated at each site with cooperator proportion *p*_0_.

### Changing environments

For some simulation runs, the metapopulation experiences a series of distinct stressful conditions. The three effects described in the previous section are applied periodically at some defined interval. Thus, when new stressful conditions are experienced, any fitness effects associated with adaptation to previous stress are removed.

### Removing population structure

Population structure can be removed by tracking a single site (a well-mixed population). To control for population size between such “unstructured” runs and the metapopulation runs described above, we let the size of the single population after growth be *S*(*p*) = *N*^2^(*S*_min_ + (*S*_max_ − *S*_min_)*p*). With the exception of the migration step (which is now absent), all other steps in the simulation cycle proceed as above.

### Parameter values, source code, and software environment

Model parameters and their values are listed in Table 1. The simulation software and all configurations are available online^1^. Simulations used Python 2.7.3, NumPy 1.9.0, and NetworkX 1.9.1 (Hagberg et al. 2008). Data analyses were performed with R 3.1.2 (R Core Team 2014).

**Table 1:** Parameters and Baseline Values.

| Symbol | Parameter Description | Value |
| --- | --- | --- |
| $N^2$ | Number of Metapopulation Sites | 625 |
| $L$ | Number of Stress Loci | 8 |
| $w_{\min}$ | Minimum Fitness Effect Per Locus | 0.45 |
| $w_{\max}$ | Maximum Fitness Effect Per Locus | 0.55 |
| $c$ | Cost of Cooperation | 0.1 |
| $z$ | Baseline Fitness | 1 |
| $T$ | Number of Simulation Cycles | 3000 |
| $S_{\min}$ | Minimum Population Size | 800 |
| $S_{\max}$ | Maximum Population Size | 2000 |
| $\mu_s$ | Mutation Rate (Stress Loci) | $10^{-5}$ |
| $\mu_c$ | Mutation Rate (Cooperation Locus) | $10^{-5}$ |
| $\mu_t$ | Mutation Rate (Tolerance to New Stress) | $10^{-5}$ |
| $m$ | Migration Rate | 0.05 |
| $d$ | Dilution Factor | 0.1 |
| $p_0$ | Initial Cooperator Proportion | 0.5 |

## Results

The major components of our hypothetical process are (1) stressful conditions create opportunities for adaptation, and (2) cooperators have greater chances to adapt due to the relatively higher reproduction that occurs in a spatially structured population. To illustrate the importance of these components, we begin by exploring the evolution of cooperation when stress adaptation and spatial structure are present or absent.

Without the opportunity for stress adaptation (*L* = 0), defectors fix rapidly (Figs. 1A and 1C). A structured population does have a slight initial lift in cooperator proportion (Fig. 1C). This pattern occurs because stress thins the metapopulation, leading to isolated populations of either cooperators or defectors. The initial lift in cooperator proportion is due to the greater productivity of cooperator populations compared to defectors. However, once migration mixes these populations, cooperators are outcompeted by defectors due to the cost of cooperation. Thus, without the possibility of stress adaptation, spatial structure does not make a big difference to the dynamics— cooperators experience rapid extinction in both cases.

Without spatial structure, defectors also have immediate advantages (Figs. 1A and 1B). However, when organisms can adapt to stress (*L* = 8) in a well-mixed population, cooperators decline less rapidly (Fig. 1B). The variance in Figure 1B reflects the fact that in some replicate runs, cooperators happen to gain the first beneficial mutation, while in other runs, defectors are first. Because the benefits of cooperation are experienced equally by both types in a well-mixed population and cooperators incur a cost, defectors actually have a greater chance of adapting (which is why the proportion of cooperators falls from its initial value over the first 1000 cycles). Cooperators in all replicates go extinct eventually; in the cases where cooperators adapt first, the rise of de novo (adapted) defectors leads to cooperator extinction.

**Figure 1:**
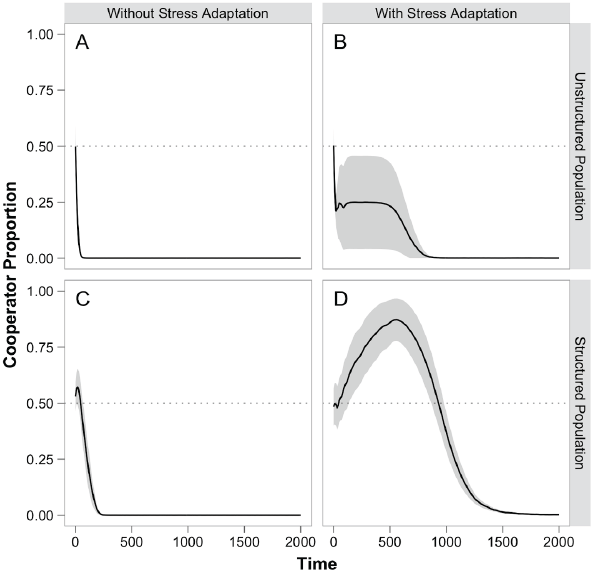
The evolution of cooperation in our model. The average proportion of cooperators across 20 replicate runs is given by the black trajectory, and shaded regions indicate 95% confidence intervals. For all model parameters not specified, the base values listed in Table 1 were used. (**A**) When there is no opportunity for adaptation to the stressful conditions (i.e., *L*, the number of stress loci, is zero) and the population is well mixed, cooperators rapidly go extinct. (**B**) However, if adaptation to the stress can occur (*L* = 8), cooperators fare better. Indeed, cooperators increase dramatically in approximately 25% of the replicate runs (before eventual extinction), whereas defectors fix quickly in the remaining set (the large variation indicates the disparity in simulation outcomes). (**C**) Without stress adaptation in a structured metapopulation, cooperators crash to extinction as in part A. (**D**) However, if adaptation to the stress is possible (*L* = 8) in a structured metapopulation, cooperators reach high proportions transiently before eventually going extinct.

When there is both the opportunity to adapt to stress and spatial structure, a dramatically different picture emerges (Fig. 1D). The greater productivity of the isolated cooperator populations creates more mutational opportunities, enabling a faster rate of adaptation to the stress. When cooperators and defectors meet through migration, the fitter cooperators can now competitively displace the defectors despite the cost of cooperation. More generally, cooperator populations are epicenters of rapid adaptation spreading and displacing defector-dominated populations. Although cooperators rise to high proportions throughout the metapopulation, this increase is ultimately transient. Because the number of stress loci is finite, cooperators will eventually discover the genotype that is most adapted to the stress. At this point, any de novo defectors will be equally stress-adapted but will save on the cost of cooperation, and thereafter displace cooperators.

Given the transient nature of cooperator success, we use the area under the cooperator proportion curve (from *T* = 0 to *T* = 3000) as a measure of cumulative cooperator presence. As cooperators spend more time at high proportions, this integral increases (Fig. 2A). By this metric, cooperators fare better as the number of stress loci increases (Fig. 2B). Not surprisingly, cooperator presence also increases as the benefit of cooperation increases (Fig. 2C) or as the cost decreases (Fig. 2D).

**Figure 2:**
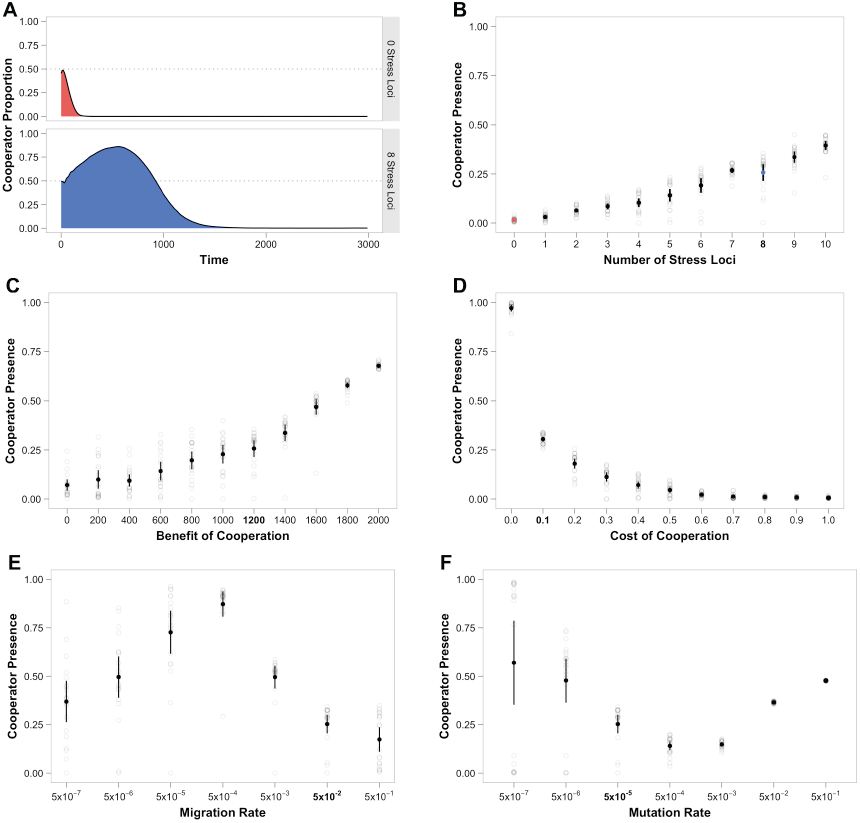
Cooperator presence as a function of various model parameters. (**A**) The area under the cooperator proportion curve is used as a measure of “cooperator presence.” The average proportions are shown in black. Thus, the average presence with *L* = 0 and *L* = 8 is given by the red and blue area, respectively. In all the graphs that follow, cooperator presence for each replicate is shown as an unfilled circle (the red and blue circles in part B correspond to the areas in part A). Average cooperator presence across 20 replicates (unfilled circles) is given by the filled circles, and bars indicate 95% confidence intervals. In each of the following graphs, one model parameter is varied, while all other parameters are set to the base values in Table 1. These base values are shown in bold text on the x axes. Cooperator presence as a function of (**B**) the number of stress loci, *L*, (**C**) the benefit of cooperation, *S*_max_ – *S*_min_, (**D**) the cost of cooperation, *c*, (**E**) the migration rate, *m*, and (**F**) the mutation rate, where we covary the mutation rate at stress loci and the cooperation locus simultaneously (*μ*_*s*_ = *μ*_*c*_).

Matters are more complicated with the rate of migration, where cooperator presence peaks at intermediate levels (Fig. 2E). At high migration rates, cooperators have insufficient time to adapt to the stress before mixing with defectors. Conversely, with low rates of migration, there is sufficient time to adapt to stress but insufficient export of adapted cooperators into less adapted defector populations.

The effects of mutation rate are also complex (Fig. 2F). At low mutation rates, we see high variance in cooperator presence—in some cases cooperators fare well (if they are the first to discover a stress mutation) while in others they do poorly (if they are not). As mutation rates increase from low levels, cooperator presence decreases. There are two contributing factors to this pattern. First, de novo defectors are more likely to arise with higher rates of mutation at the cooperation locus, *μ*_*c*_. Second, the isolated defector populations are better able to adapt to the stress with higher rates of mutation at the stress loci, *μ*_*s*_. As mutation rates continue to increase, cooperator presence increases due to a higher cooperator proportion at mutation-selection balance (see Supplemental Figure S1).

For all of the results above, the metapopulation adapts in response to a single stressful environment. If the metapopulation instead faces a series of stressful environments, we see that cooperator proportion can be maintained at high values (Fig. 3A). However, the frequency at which environmental changes occur is crucial. This frequency must be greater than a critical value in order for cooperators to avoid extinction (Fig. 3B). As the frequency of change increases above this value, a fresh round of adaptation salvages an otherwise doomed cooperator lineage. At the highest frequencies of environmental change, there is not sufficient time for adaptation to any new stress (given the rapid barrage of changing harsh conditions). Cooperators fare well in these quickly changing environments solely due to the continual thinning effect associated with stress and the ensuing positive assortment (see Supplemental Figure S2).

**Figure 3:**
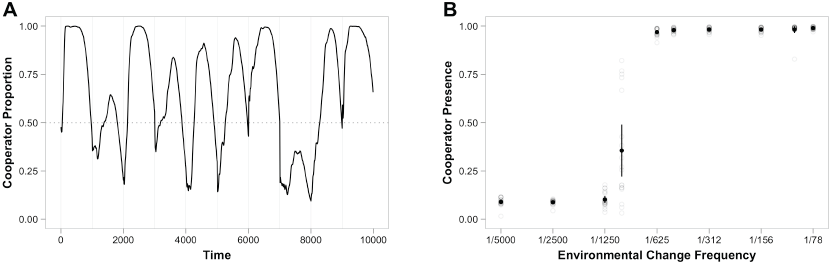
Evolution of cooperation in changing environments. (**A**) When new stressful conditions occur every 1000 cycles (faint vertical lines), cooperators remain at high proportion for long periods of time (base parameter values from Table 1 are used here). (**B**) The frequency of environmental change must be sufficiently large (greater than ∼1/1100) for cooperators to have a sustained presence. Replicate runs are given by unfilled circles, filled black circles display the average across replicates, and bars indicate 95% confidence intervals.

## Discussion

In our model, cooperation evolves through a form of hitchhiking. In order for this costly trait to hitchhike, there must be “rides” available; that is, there must be opportunities for beneficial mutations. In our simulated scenario, a stressful environment provides such opportunities for adaptation. However, evolution by the Hankshaw effect involves more than simple hitchhiking; rather, it requires that cooperators have a greater chance of catching a ride than defectors, which can occur if cooperators have more reproductive opportunities. In our simulation, the combination of stress-induced thinning and spatially limited dispersal produces positive assortment and higher cooperator productivity follows. With opportunities to adapt to stress and with limited dispersal, cooperators can experience sustained increases in frequency (Figure 1).

In the process we have outlined, the cooperation allele is a hitchhiking passenger with a “driving” allele that confers stress adaptation. While the pace of stress adaptation depends on social traits, the stress adaptation itself is inherently non-social. The idea that adaptation to non-social aspects of the environment can affect social evolution has been explored both empirically (Waite and Shou 2012, Morgan et al., 2012, Asfahl et al. 2015) and theoretically (Quigley et al., 2012). These previous studies focus on well-mixed populations of cooperators and defectors, where a beneficial mutation can arise in either a cooperator or a defector. If this mutation occurs in the cooperator background, and if the benefit of this mutation outweighs the cost of cooperation, then the cooperators may displace the defectors. This process has been termed an “adaptive race” in the sense that the cooperator and defector are in a race for the first adaptive mutation (Waite and Shou, 2012). In the adaptive race, the cooperation allele does not have a greater chance than the defection allele of hitchhiking. Consequently, the average proportion of cooperators (e.g., across many replicate populations) is not expected to increase (see Fig. 1B). In contrast, in a spatially structured population, the cooperation allele directly increases its probability of catching a lift, leading to an actual increase in the average cooperator proportion (see Fig. 1D).

Since Hamilton’s pioneering work (Hamilton 1963, 1964), it is well known that any mechanism by which cooperators cluster together facilitates the evolution of cooperation (Hamilton 1975, Wilson 1975, Pepper and Smuts 2002, West et al. 2007b, Fletcher and Doebeli 2009, Nadell et al. 2010). In our model, restricted dispersal within the metapopulation after stress-induced thinning leads to a high degree of clustering. Such positive assortment alone does give an initial boost to cooperators, even without adaptation to stress (compare Fig. 1C to 1A). However, the effect of spatial clustering is much more profound when stress adaptation is possible (compare Fig. 1D to 1B). In the Supplement, we demonstrate that the combination of stress-adaptation and limited dispersal (but *without* stress-induced thinning) allows for high cooperator success (Supplemental Figure S3). Overall, we see that limited dispersal is an important, but not alone sufficient, condition for the evolution of cooperation by the Hankshaw effect.

While the operation of our process in natural systems is an empirical issue, our model suggests that the evolution of cooperation by the Hankshaw effect can occur under a broad set of conditions. Indeed, the basic assumptions of the model are fundamental features of many biological populations. For instance, cooperation in the form of the production of public goods or competitive restraint in the use of common resources has been shown to increase population size (Kerr et al. 2006, Diggle et al. 2007, Eshelman et al. 2010, Xavier et al. 2011, Drescher et al. 2014), a critical assumption of our model. Moreover, many populations are naturally structured, from microbial biofilms to passively dispersed plants to sessile or territorial animals (Hutchings 1997, Tilman and Karieva 1997, Nadell et al. 2009). Finally, in natural systems, harsh environmental conditions can impact the survival and reproduction of organisms. In fact, the edges of a species’ range can sometimes be defined by stressful conditions (Connell 1961, Sexton et al. 2009, Hargreaves et al. 2014). It is likely that these assumptions will be simultaneously satisfied in a natural context.

Cooperation is not the only kind of trait that can evolve by the Hankshaw effect. It is possible that spiteful traits (i.e., phenotypes that harm others at a personal cost; Hamilton 1970, Gardner and West 2006) may also create more opportunities to hitchhike within structured populations. For example, the production of toxins in bacteria (bacteriocins; Chao and Levin 1981, Riley and Wertz 2002, Kerr et al. 2002, Inglis et al. 2009) or the possession of features that enhance flammability in plants (Mutch 1970, Williamson and Black 1981, Schwilk 2003) are spiteful traits that could evolve through the Hankshaw effect. Specifically, individuals with these traits create empty patches in their population at an extreme personal cost (by lysing or burning); and adaptation by relatives (clone mates or offspring) may occur at a higher rate (see Schwilk and Kerr 2002 for a full treatment).

In our model, the success of costly social traits (cooperation or spite) is merely transient when de novo defectors can arise and stress adaptation is limited. However, these social traits can experience more than transient success if the environment changes continually (Figure 3). Periodic change (e.g., diurnal or seasonal cycles) is experienced by many biological populations (Fretwell 1972, McClung 2006). Additionally, organisms themselves can alter their environment, a process termed “niche construction” (Odling-Smee et al. 2003, Laland et al. 1999). Thus, cycles of environmental change can be generated by the evolving system. For instance, suppose that as population density increases, new stressful conditions become more likely (e.g., at high population density, new pathogens may be more likely to spread, creating new selective pressures for the host population). In such a case, cooperation may be maintained at high proportion indefinitely.

In summary, we have explored a scenario where an allele improves its own prospects for hitchhiking. While the most straightforward case involves direct effects of the allele on its owner (e.g., a mutator allele), here we have explored a more subtle case. Specifically, the increased probability of hitchhiking of our focal allele occurs due to its social impact inside a structured population. In the process, the social behavior increases in proportion despite its costs. Common explanations for the evolution of costly social traits (genetic hitchhiking and positive assortment) are elements of our model. However, the effect of bringing these elements together has remained unexplored before this study. Our theoretical results reveal this unification to be synergistic. Given the biological plausibility of our theoretical assumptions, the Hankshaw effect will be an interesting focus for the study of social evolution in the future.

## Acknowledgements

We thank P. Conlin, C. Lee, W. Shou, A. Waite, and the ShoKer and SQuEE groups for discussion during the early stages of this work. We thank J. Cooper, S. Estrela, K. Foster, C. Glenney, H. Jordt, W. Shou, S. Singhal, A. Titus, K. van Raay, and L. Zaman for helpful comments on the manuscript. We are particularly grateful to Pete Hammarlund for simulation advice, as well as for raising a concern about environmental change and his legwork in finding apparent links to this topic in previous work. This material is based upon research supported by the National Science Foundation under Grant 1309318 (Postdoctoral Research Fellowship in Biology to BDC), Cooperative Agreement DBI-0939454 (BEACON STC) and Grant DEB0952825 (CAREER Award to BK). Computational resources were provided by an award from Google (to BDC and BK).

## Footnotes

1 Brian Connelly. 2015. Model for The Evolution of Cooperation by the Hankshaw Effect. Zenodo. 10.5281/zenodo.16490.

